# Conservation of a gene cluster reveals novel cercosporin biosynthetic mechanisms and extends production to the genus *Colletotrichum*

**DOI:** 10.1101/100545

**Authors:** Ronnie de Jonge, Malaika K. Ebert, Callie R. Huitt-Roehl, Paramita Pal, Jeffrey C. Suttle, Rebecca E. Spanner, Jonathan D. Neubauer, Wayne M. Jurick, Karina A. Stott, Gary A. Secor, Bart P.H.J. Thomma, Yves Van de Peer, Craig A. Townsend, Melvin D. Bolton

**Affiliations:** Department of Plant Systems Biology, VIB, Ghent, Belgium.; Department of Plant Biotechnology and Bioinformatics, Ghent University, Ghent, Belgium.; Bioinformatics Institute Ghent, Ghent University, B9052 Gent, Belgium.; Plant-Microbe Interactions, Department of Biology, Faculty of Science, Utrecht University, Utrecht, The Netherlands.; Northern Crop Science Laboratory, United States Department of Agriculture, Fargo, ND, United States.; Department of Plant Pathology, North Dakota State University, Fargo, ND, United States.; Laboratory of Phytopathology, Wageningen University, Wageningen, the Netherlands.; Department of Chemistry, The Johns Hopkins University, Baltimore, MD, United States.; Food Quality Laboratory, United States Department of Agriculture, Beltsville, MD, United States.; Department of Genetics, Genomics Research Institute, University of Pretoria, Pretoria, South Africa.

**Keywords:** natural product, perylenequinone, secondary metabolite, cercosporin, *Cercospora*, *Colletotrichum*

## Abstract

Species in the genus *Cercospora* cause economically devastating diseases in sugar beet, maize, rice, soy bean and other major food crops. Here we sequenced the genome of the sugar beet pathogen *C. beticola* and found it encodes 63 putative secondary metabolite gene clusters, including the cercosporin toxin biosynthesis (*CTB*) cluster. We show that the *CTB* gene cluster has experienced multiple duplications and horizontal transfers across a spectrum of plant pathogenic fungi, including the wide host range *Colletotrichum* genus as well as the rice pathogen *Magnaporthe oryzae*. Although cercosporin biosynthesis has been thought to-date to rely on an eight gene *CTB* cluster, our phylogenomic analysis revealed gene collinearity adjacent to the established cluster in all *CTB* cluster-harboring species. We demonstrate that the *CTB* cluster is larger than previously recognized and includes *cercosporin facilitator protein* (*CFP*) previously shown to be involved with cercosporin auto-resistance, and four additional genes required for cercosporin biosynthesis including the final pathway enzymes that install the unusual cercosporin methylenedioxy bridge. Finally, we demonstrate production of cercosporin by *Colletotrichum fioriniae*, the first known cercosporin producer within this agriculturally important genus. Thus, our results provide new insight into the intricate evolution and biology of a toxin critical to agriculture and broaden the production of cercosporin to another fungal genus containing many plant pathogens of important crops worldwide.

**Significance Statement:** Species in the fungal genus *Cercospora* cause diseases in many important crops worldwide. Their success as pathogens is largely due to the secretion of cercosporin during infection. We report that the cercosporin toxin biosynthesis (*CTB*) cluster is ancient and was horizontally transferred to diverse fungal pathogens on an unprecedented scale. Since these analyses revealed genes adjacent to the established *CTB* cluster, we evaluated their role in C. beticola to show that four are necessary for cercosporin biosynthesis. Finally, we confirmed that the apple pathogen *Colletotrichum fioriniae* produces cercosporin, the first case outside the family Mycosphaerellaceae. Other *Colletotrichum* plant pathogens also harbor the *CTB* cluster, which points to a wider concern that this toxin may play in virulence and human health.

---

*Cercospora* are among the most speciose genera in all Fungi. First described in 1863 (1), the genus has sustained a long history, largely due to notoriety as the causal agent of leaf spot diseases in a wide range of plants including agriculturally important crops such as sugar beet, soybean, maize and rice that together account for hundreds of millions of dollars in lost revenue annually to growers worldwide. Although *Cercospora* spp. share several characteristics associated with pathogenicity, such as penetration through natural openings and extracellular growth during the biotrophic stage of infection, most rely on the production of the secondary metabolite (SM) cercosporin to facilitate infection (2, 3). Studies spanning nearly 60 years have made cercosporin a model perylenequinone (4), a class of SMs characterized by a core pentacyclic conjugated chromophore that gives rise to its photoactivity. When exposed to ambient light, cercosporin is a potent producer of reactive oxygen species in the presence of oxygen (5) with a quantum efficiency of >80% (6). This small molecule is lipophilic and can readily penetrate plant leaves leading to indiscriminate cellular damage within minutes of exposure (7). Indeed, cercosporin is nearly universally toxic to a wide array of organisms including bacteria, mammals, plants and most fungal species with the key exception of cercosporin-producing fungi, which exhibit cercosporin auto-resistance. To our knowledge, cercosporin is only known to be produced by *Cercospora* spp. with the single exception of the brassica pathogen *Pseudocercosporella capsellae* (8). However, *Pseudocercosporella* and *Cercospora* are phylogenetically closely-related, residing in a large clade within the Mycosphaerellaceae (9).

In contrast to the large body of information on cercosporin biology spanning several decades (10, 11), the cercosporin toxin biosynthesis (*CTB*) gene cluster was only recently resolved in *C. nicotianae* (12). The keystone enzyme for cercosporin biosynthesis, CTB1, bears all the hallmarks of an iterative, non-reducing polyketide synthase (NR-PKS) (13). Using *CTB1* as a point of reference, the complete *C. nicotianae CTB* gene cluster was determined to consist of eight contiguous genes of which six are believed to be responsible for cercosporin assembly (*CTB1*, *2*, *3*, *5*, *6*, and *7*) (12, 14). The zinc finger transcription factor CTB8 co-regulates expression of the cluster (12), while the major facilitator superfamily (MFS) transporter CTB4 exports the final metabolite (15). Downstream of the *CTB* cluster are two open reading frames (ORFs) encoding truncated transcription factors, while loci designated as *ORF9* and *ORF10* upstream of the *CTB* cluster are not regulated by light and are not hypothesized to encode proteins with metabolic functions (12). Consequently, the clustering of eight genes with demonstrated co-regulation by light that are flanked by ORFs with no apparent role in cercosporin biosynthesis has suggested that cercosporin production relies on the eight-gene *CTB* cluster (12). In this study, we used an evolutionary comparative genomics approach to show that the *CTB* gene cluster underwent multiple duplication events and was transferred horizontally across large taxonomic distances. Since these horizontal transfer events included genes adjacent to the canonical eight gene *CTB* cluster, we used reverse genetics to show that the *CTB* cluster includes additional genes in *C. beticola*, including one gene that was previously shown to be involved with cercosporin auto-resistance (16) and four previously unrecognized genes involved with biosynthesis. The *CTB* cluster was found in several *Colletotrichum* (*Co*.) species, and we confirmed that the apple pathogen *Co. fioriniae* can also produce cercosporin. As all earlier understanding of cercosporin biosynthesis has been unwittingly limited by a truncated set of genes in *Cercospora* spp., the full dimension of the gene cluster provides deeper insight into the evolution and dissemination of a fungal toxin critical to world-wide agriculture.

## Results

### Secondary metabolite cluster expansion in *Cercospora beticola*

*C. beticola* strain 09-40 was sequenced to 100-fold coverage and scaffolded with optical and genome maps, resulting in 96.5% of the 37.06 Mbp assembly being placed in 12 supercontigs of which 10 are assumed to be chromosomes. Despite their ubiquitous presence in nature and cropping systems, genome sequences of *Cercospora* spp. are not well-represented in public databases. Therefore, to aid comparative analysis within the *Cercospora* genus we also sequenced the genome of *C. berteroae* and reassembled the genome of *C. canescens* (17) (*SI Appendix*, Table S1). To identify gene clusters responsible for biosynthesis of aromatic polyketides in *C. beticola*, we mined the genome to identify all SM clusters (18) and compared these with predicted clusters in related Dothideomycete fungi. The *C. beticola* genome possesses a total of 63 predicted SM clusters of several classes (Table S2), representing a greatly expanded SM repertoire compared to closely related Dothideomycetes (*SI Appendix*, Table S3). To identify the *C. beticola* PKS cluster responsible for cercosporin biosynthesis, we compared the sequence of the *C. nicotianae CTB* cluster (12) with predicted PKS clusters of *C. beticola*. The *C. beticola* PKS *CBET3_00833* (*CbCTB1*) and flanking genes (*CBET3_00830* – *CBET3_00837*) were ~96% identical to *C. nicotianae CTB1 – CTB8* and all genes were collinear, strongly suggesting this region houses the *CTB* cluster in *C. beticola*.

### Repeated duplication and lateral transfer of the cercosporin biosynthetic cluster

To study the evolutionary relationships *of C. beticola* PKSs, we conducted large-scale phylogenomic analyses that included various previously characterized PKSs from selected species (19). Since resolving orthologous relationships among PKSs can predict the type of SM that will be synthesized, we first built a phylogenetic tree of the conserved core β-ketoacyl synthase (KS) domains of each PKS that resulted in separating PKS enzymes into four major groups (*SI Appendix*, Fig. S1*A*). Among the eight *C. beticola* NR-PKSs, phylogenetic analysis revealed significant similarity between *CbCTB1*, *CBET3_10910-RA*, and *CBET3_11350-RA* which cluster at the base of the cercosporin clade (*SI Appendix*, Fig. S1*B*). Interestingly, genes flanking *CBET3_10910-RA*, but not *CBET3_11350-RA*, were also strikingly similar to *CbCTB* cluster genes (Fig. 1). Consequently, we hypothesize that the *CBET3_10910* SM cluster is the result of a *CTB* cluster duplication. Since duplicated SM gene clusters appeared to be relatively rare in fungi (20), we investigated the origin and specificity of the *CTB* cluster and the putative duplication by searching for *Cb*CTB1 homologs against a selected set of 48 published Ascomycete proteomes (*SI Appendix*, Table S4) representing a diverse group of fungal orders. We identified *Cb*CTB1 orthologs in *Cercospora* spp. *C. berteroae* and *C. canescens* and confirmed its presence in *Cladosporium fulvum* (19) and *Parastagonospora nodorum* (21). Surprisingly, seven additional orthologs were identified in Sordariomycete species *Co. orbiculare*, *Co. gloeosporioides*, *Co. fioriniae*, *Co. graminicola*, *Co. higginsianum*, and *Magnaporthe oryzae* as well as one in the Leotiomycete *Sclerotinia sclerotiorum* (*SI Appendix*, Fig. S2*A*), representing diverse taxa harboring *CTB1*. Analysis of sequence similarity showed that intra-species (*Cb*CTB1 - CBET3_10910-RA) sequence identity (45%) was lower than the inter-species identity (e.g. *Cb*CTB1 and *C. fulvum* CTB1 (Clafu1_196875) are 55% similar; *SI Appendix*, Table S5), suggesting that the *CTB1* duplication event was ancient and occurred prior to Dothideomycete speciation.

**Figure 1.**
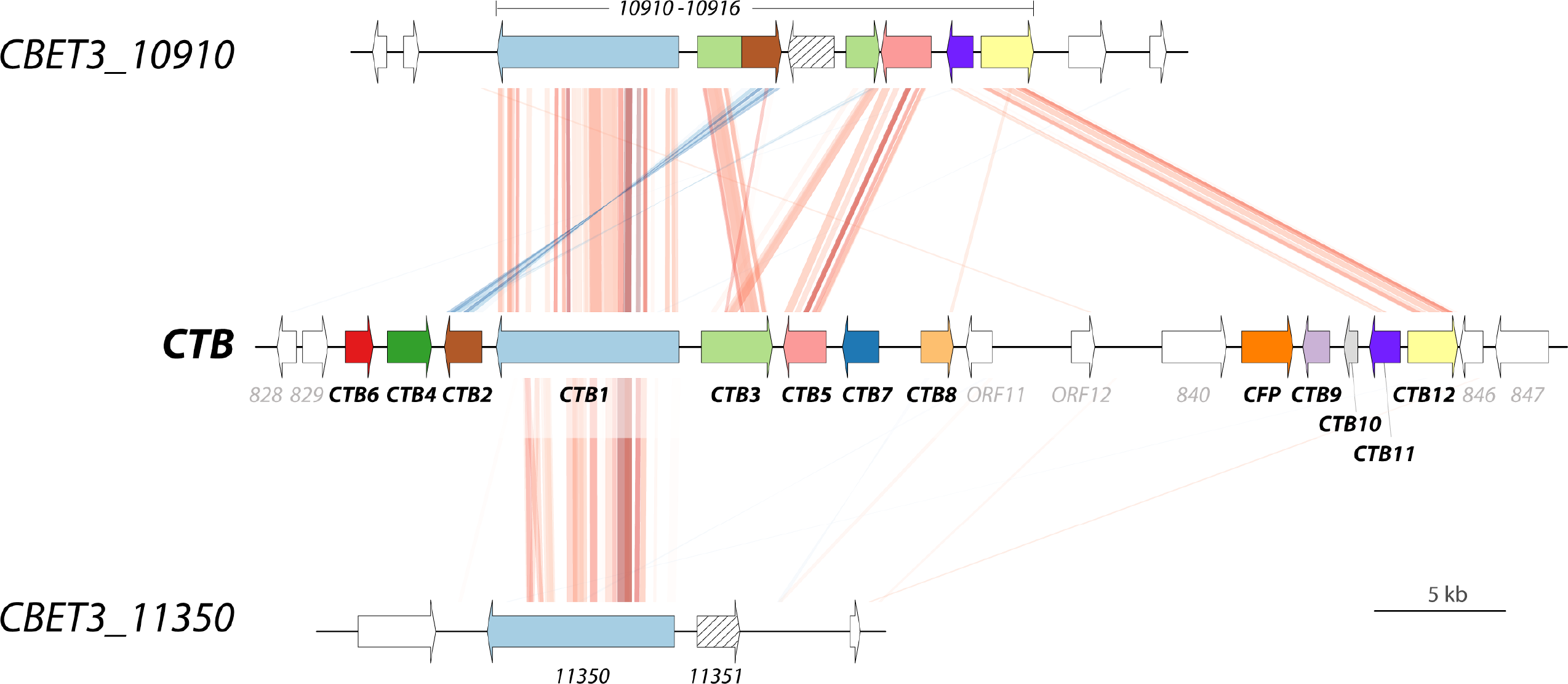
The cercosporin biosynthetic cluster is duplicated and maintained in C. beticola. *CBET3_10910* and flanking genes are syntenic with the *CTB* cluster (*CBET3_00833* and flanking genes) in *C. beticola*. Alignment lines correspond to DNA fragments exhibiting significant similarity when the genomic regions comprising the gene clusters are compared with tBLASTx. Direct hits are displayed in red, whereas complementary hits are in blue. The intensity of the alignments represents the percentage similarity ranging from 23 to 100 percent. Genes flanking *CBET3_11350-RA* were not syntenic with *CTB* cluster genes.

To develop a ‘phylogenetic roadmap’ that may explain *CTB1* evolution, we used the process of ‘reconciliation’ that takes into account both species and gene histories (22). Although not conclusive, reconciliation considers the costs of evolutionary events (i.e. gene duplications, transfers, and/or losses) to explain the most parsimonious evolutionary route to the present scenario (23). Reconciliation of the species tree (*SI Appendix*, Fig. S3) with the CTB1 protein tree revealed that the predicted evolutionary history of CTB1 can be characterized by four duplications, three transfers and wide-spread loss to most species analyzed (*SI Appendix*, Fig. S4*A*), and further corroborates our hypothesis that the *CTB1* duplication event (D1) occurred prior to Dothideomycete speciation. Reconciliation revealed an ancient transfer in which the lineage leading to *S. sclerotiorum* acquired the duplicated *CTB1* from the last common ancestor of *Cercospora* spp. (T1; *SI Appendix*, Fig. S4*A*). Duplications 2-4 (D2-4) arose after lateral transfer (T2) of *CTB1* into the last common ancestor of the *Glomerellales*. *CTB1* was then transferred (T3) from a common ancestor in the *Glomerellales* to *M. oryzae* (*SI Appendix*, Fig. S4*A*).

We extended the search for CTB cluster protein orthologs by scanning the 48 proteomes for homologs of *Cb*CTB2 (CBET3_00830) to *Cb*CTB8 (CBET3_00837) followed by phylogenetic tree construction and subtree selection (*SI Appendix*, Fig. S2). This resulted in the identification of orthologs in the same set of species previously listed to contain *CTB1*, with the only exceptions in cases where *CTB* gene homologs were lost in a species. Although the loss of CTB6 and CTB7 orthologs limits reconciliation analysis of these gene families, reconciliation of the subtrees for CTB2, CTB3, CTB4, CTB5 and CTB8 (*SI Appendix*, Fig. S4) supported a similar scenario as proposed for CTB1, which is summarized as two duplications and two horizontal transfer events that explain the present-day *CTB* scenario (Fig. 2). However, a slightly less parsimonious explanation is a single transfer to an ancestral Glomerellales species, followed by wide-spread loss in most species in this lineage except for *M. oryzae* and the analyzed *Colletotrichum* spp. (Fig. 2; *SI Appendix*, Table S6).

**Figure 2.**
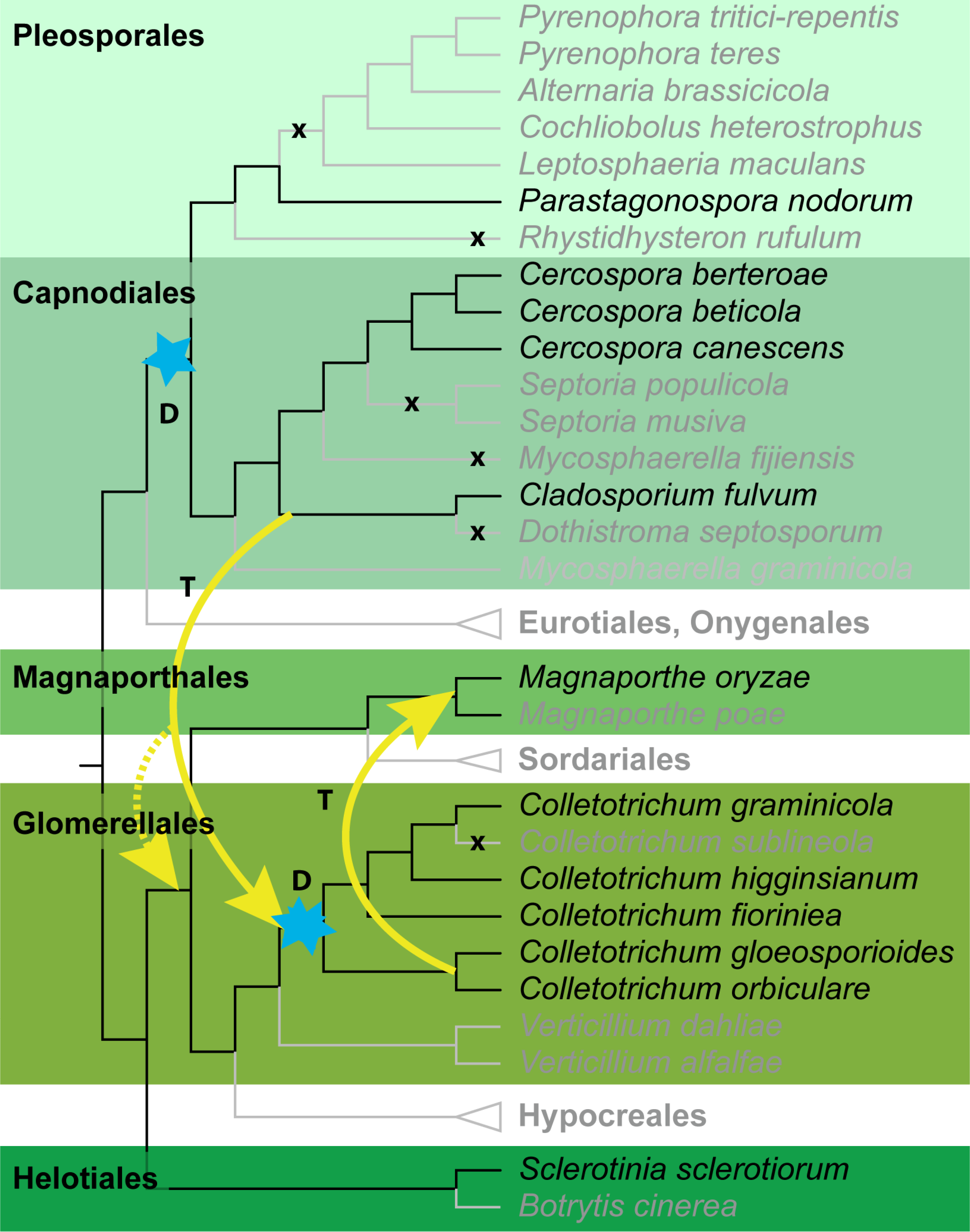
Phylogeny of *Cercospora* spp. and related Ascomycete fungi. Cladogram showing the phylogenetic relationship of *Cercospora* spp. and 45 other sequenced fungi. The unscaled tree was constructed using CVTree. Duplication nodes are marked with blue stars, losses are indicated by the crosses and transfers are highlighted by yellow arrows. Species without the *CTB* cluster are depicted in grey, those encompassing it are in black. The alternative and slightly less parsimonious scenario involving a single transfer into the last common ancestor of the Magnaporthales and the Glomerellales is shown by the dashed arrow.

### Extension of the predicted cercosporin biosynthetic cluster based on microsynteny

To further examine the *CTB* clusters across all recipient species we generated pairwise alignments relative to the *C. beticola CTB* cluster and flanks. To our surprise, we observed a striking level of similarity outside of the known eight *CTB* genes on the 3’ end of the cluster (Fig. 3) in all *CTB* containing genomes. To investigate whether the amount of microsynteny observed for *CTB* cluster and these flanking genes can be reasonably expected when comparing *Dothideomycete* and *Sordariomycete* genomes, we assessed the genome-wide microsynteny between the genomes of *C. beticola* and *Co. gloeosporioides* and *C. beticola* and *M. oryzae*. This analysis identified the *CTB* cluster together with its flanking genes as having the highest level of microsynteny among all regions in the genome between *C. beticola* and *Co. gloeosporioides*, and showed that the observed *CTB* microsynteny between *C. beticola* and *M. oryzae* was significantly higher than the genome-wide average (Fig. 4). Likewise, CTB protein similarity between *C. beticola* and *Colletotrichum* spp., and to a lesser degree with *M. oryzae*, is higher compared to the genome-wide average (*SI Appendix*, Fig. S5, S6). Thus, we hypothesized that these flanking genes are likely part of the *C. beticola CTB* cluster, and consequently it is significantly larger than previously described (12). To test this, we first determined the relative expression of all eight established *C. beticola CTB* genes as well as a number of flanking genes (*CBET3_00828* to *CBET3_00848*) under light (cercosporin-inducing) compared to dark (cercosporin-repressing) conditions, which showed that all candidate *CTB* genes on the 3’ flank were induced in the light except *CBET3_00846* and *CBET3_00848* (*SI Appendix*, Table S7). Functional annotation of these novel, induced genes revealed one non-conserved phenylalanine ammonia lyase (CBET3_00840), the cercosporin facilitator protein (CFP) (16) (CBET3_00841), a candidate α-ketoglutarate-dependent dioxygenase (CBET3_00842), an EthD-domain containing protein (CBET3_00843), a β-ig-h3 fasciclin (CBET3_00844), a laccase (CBET3_00845) and protein phosphatase 2A (CBET3_00847; *SI Appendix*, Table S7), which have functions associated with multi-domain enzymes or polyketide biosynthesis in fungi or bacteria (12, 24-29). Phylogenetic analyses of these flanking genes and reconciliation of their respective protein phylogenies (*SI Appendix*, Fig. S2) with the species tree suggest that all genes except *CBET3_00840*, *CBET3_00846*, and *CBET3_00847* have undergone highly similar evolutionary trajectories as the established *CTB* cluster genes (Fig. 2, *SI Appendix*, Fig. S4) suggesting that the *CTB* cluster was transferred as a whole at least once, followed by species-specific evolutionary trajectories involving frequent gene loss (Fig. 2) as well as gene gain.

**Figure 3.**
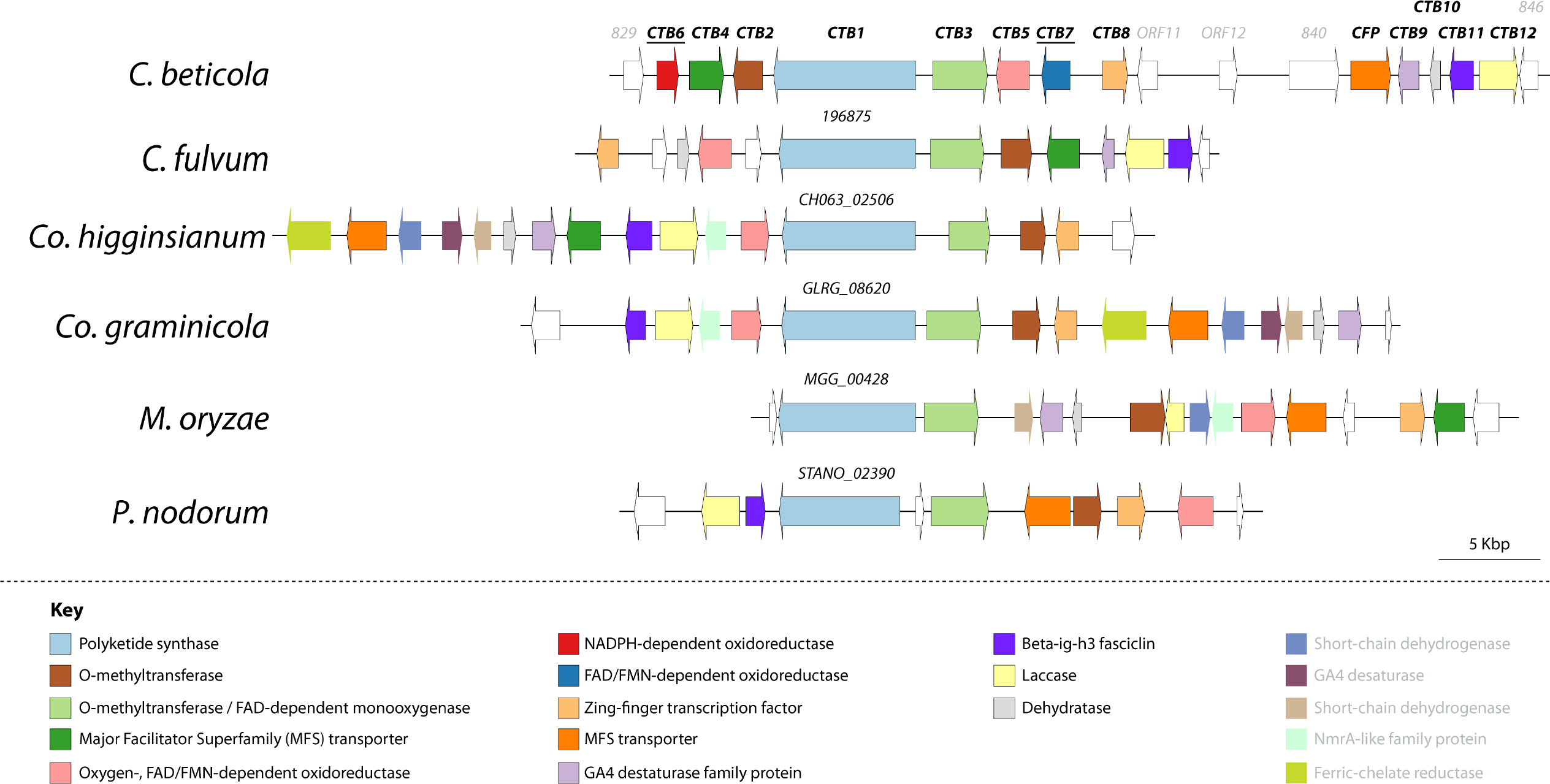
Synteny and rearrangements of the conserved *C. beticola* cercosporin biosynthetic cluster. The cercosporin biosynthetic cluster in *C. beticola* (Cb), top line, and flanking genes are conserved in *Cladosporium fulvum*, *Co. higginsianum*, *Co. graminicola*, *M. oryzae* and *Parastagonospora nodorum*. For all species the displayed identifiers are transcript IDs and the corresponding sequences can be retrieved from JGI MycoCosm or ORCAE. *CTB* orthologs are colored relative to the *C. beticola CTB* cluster genes and the color key, as well as annotated functions, are highlighted below the *CTB* cluster graphic.

**Figure 4.**
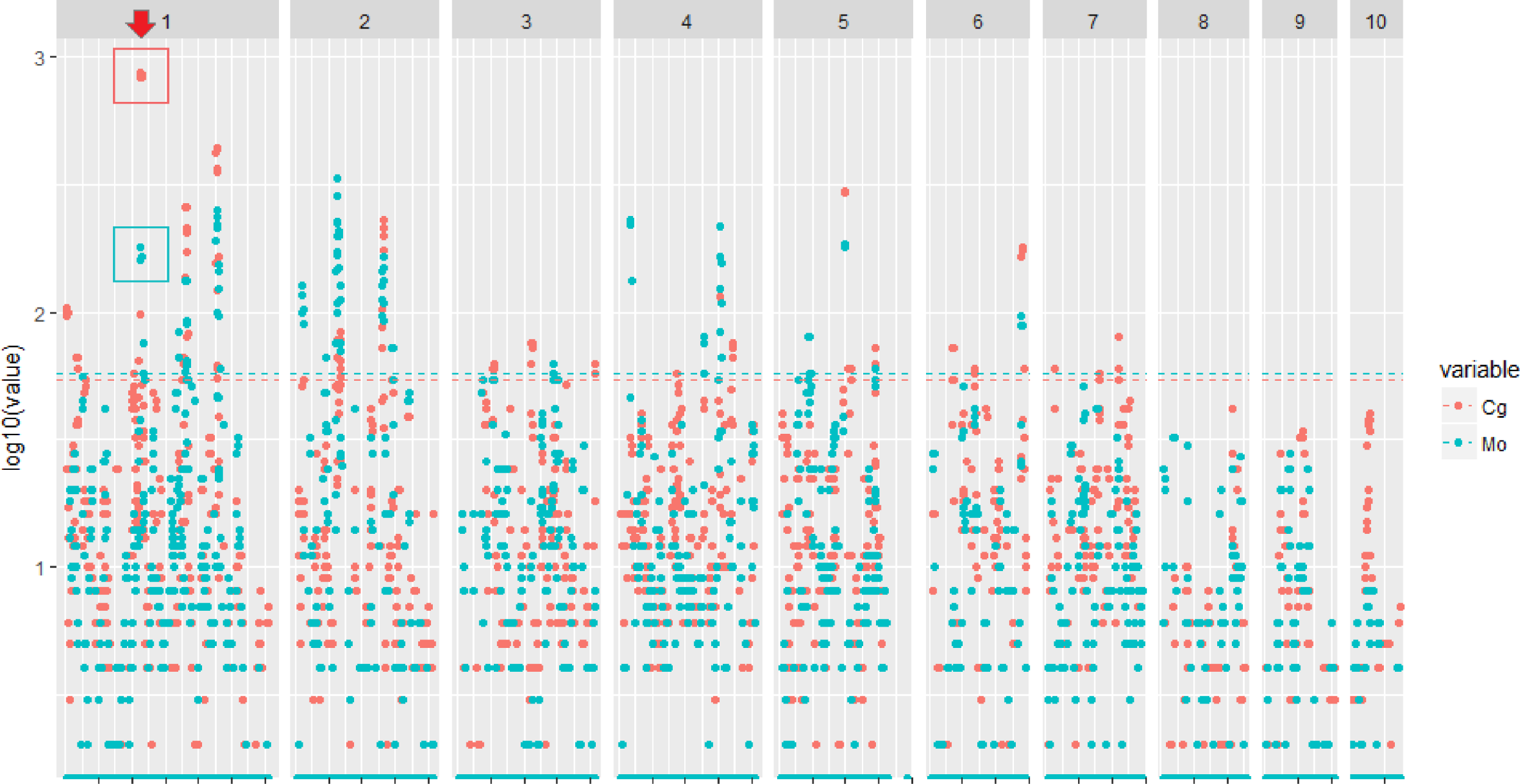
*CTB* cluster microsynteny conservation segregates from the genome-wide average. The genome-wide, gene-by-gene microsynteny between *Cercospora beticola* and *Colletotrichum gloeosporioides* (Cg, red), and between *C. beticola* and *M. oryzae* (Mo, blue), across the ten assembled *C. beticola* chromosomes is shown. Each dot represents one *C. beticola* gene and its respective microsynteny score. The red arrow indicates the position of the *CTB* cluster on chromosome 1 and coincides with high microsynteny in both *C. gloeosporioides* and *M. oryzae*. The dashed lines represent the 99^th^ quantile of the microsynteny scores for both comparisons.

### Novel CTB genes are essential for cercosporin biosynthesis

To confirm individual gene contributions for cercosporin production, we generated single gene deletion mutants of all candidate genes from *CBET3_00840* to *CBET3_00846* and tested their ability to produce cercosporin. These assays showed that cercosporin production in Δ*CBET3_00844* and Δ*CBET3_00845* mutants was abolished, while Δ*CBET3_00842* and Δ*CBET3_00843* mutants accumulated only a red, cercosporin-like metabolite that migrated differently in potato dextrose agar (PDA) culture plates and thin layer chromatography (TLC) (*SI Appendix*, Fig. S7), and exhibited a different profile obtained via high-performance liquid chromatography (HPLC) compared to cercosporin (Fig. 5). Other mutants produced compounds with HPLC profiles like cercosporin (Fig. 5), suggesting these genes are not involved with cercosporin biosynthesis. Taken together, these results corroborate our hypothesis that the *CTB* cluster extends to at least *CBET3_00845* at the 3’ side and includes four novel *CTB* biosynthetic genes as well as *CbCFP*. Consequently, we propose naming genes *CBET3_00842*, *CBET3_00843*, *CBET3_00844* and *CBET_00845* as *CbCTB9* to *CbCTB12*, respectively (*SI Appendix*, Table S7).

**Figure 5.**
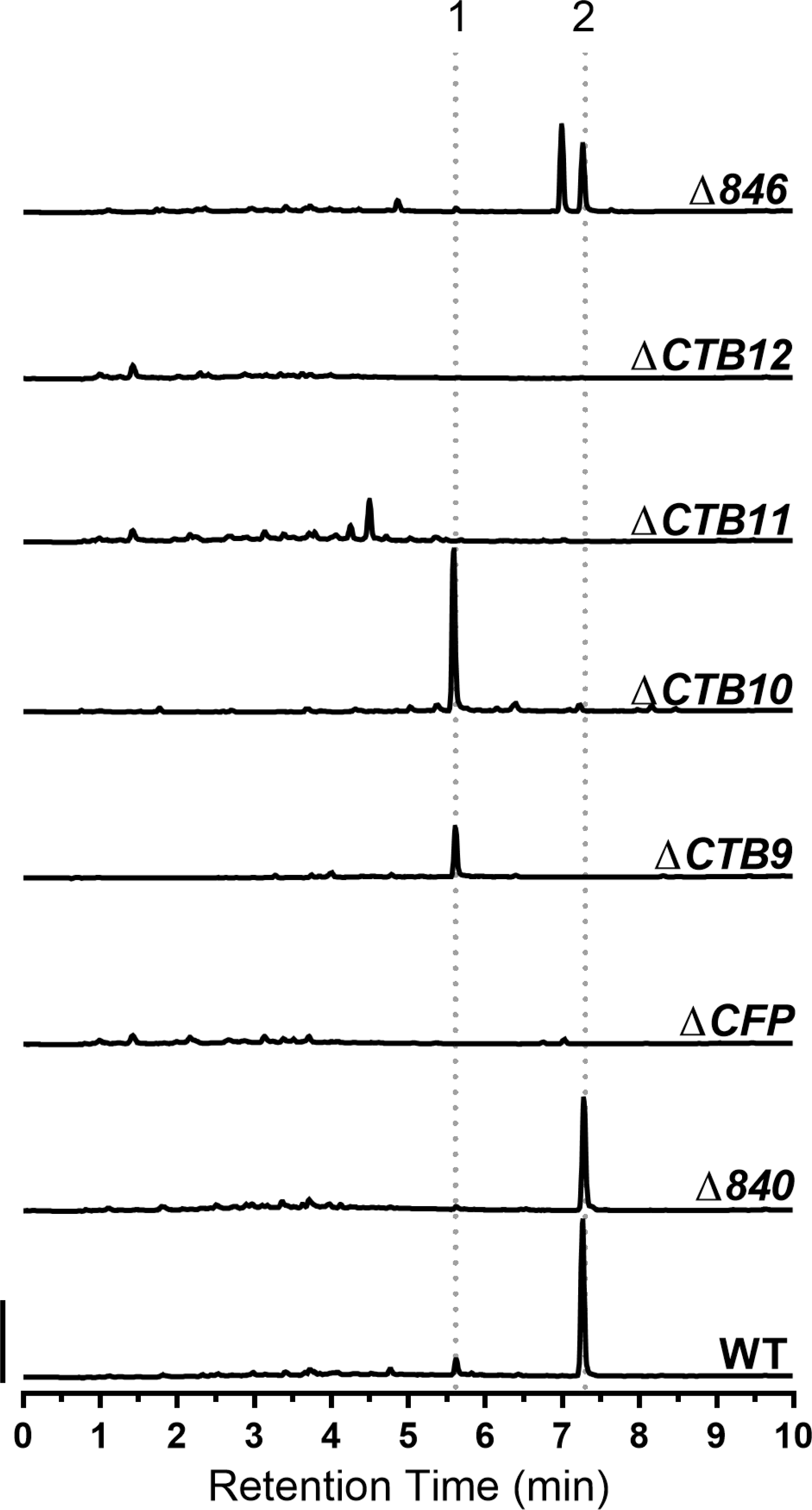
Analysis of cercosporin production in *CTB* mutants of *C. beticola*. Site-directed knock-out mutants in genes *CBET3_00840*, *CFP (CBET3_00841)*, CTB9 (*CBET3_00842*), CTB10 (*CBET3_00843*), CTB11 (*CBET3_00844*), CTB12 (*CBET3_00845*) and *CBET3_00846* were assayed for cercosporin production by HPLC. Cercosporin extracted from *C. beticola* strain 10-73-4 (WT) was used as a positive control. Pre-cercosporin (1) and cercosporin (2) are indicated by dashed lines. Scale bar indicates 250 mAu.

### Pre-cercosporin isolation and characterization

To characterize the red metabolite that accumulated in Δ*CTB9* and Δ*CTB10* mutants, an ethyl acetate extract of the collected mycelia was analyzed by reverse phase HPLC. At 280 nm, a single peak was observed in both mutant extracts with identical retention time and UV-Vis spectra (Fig. 5,6). This peak was compared to a reference sample of cercosporin produced by wild-type *C. beticola*. The retention time of this peak was shorter than that of cercosporin suggesting a more polar metabolite. Comparison of the UV-vis spectra (Fig. 6*A-C*) of the unknown compound and cercosporin revealed nearly identical chromophores, suggesting close structural relation. The exact mass of the metabolite from the mutants was determined (Δ*CTB9*: m/z = 537.1762, Δ*CTB10*: m/z = 537.1757, [M+H^+^]), consistent with the elemental composition C_29_H_28_O_10_. This mass is 2 Da greater than that of cercosporin (+2 hydrogens), which led to a proposed structure for pre-cercosporin (Fig. 6*D*). Alternative hydroquinones of cercosporin could be excluded simply on the basis of the UV-vis spectral information and chemical instability. The presence of a free phenol in pre-cercosporin in place of the unusual 7-membered methylenedioxy of cercosporin is consonant with the red shift of the long wavelength λ_max_ and the shorter HPLC retention time.

**Figure 6.**
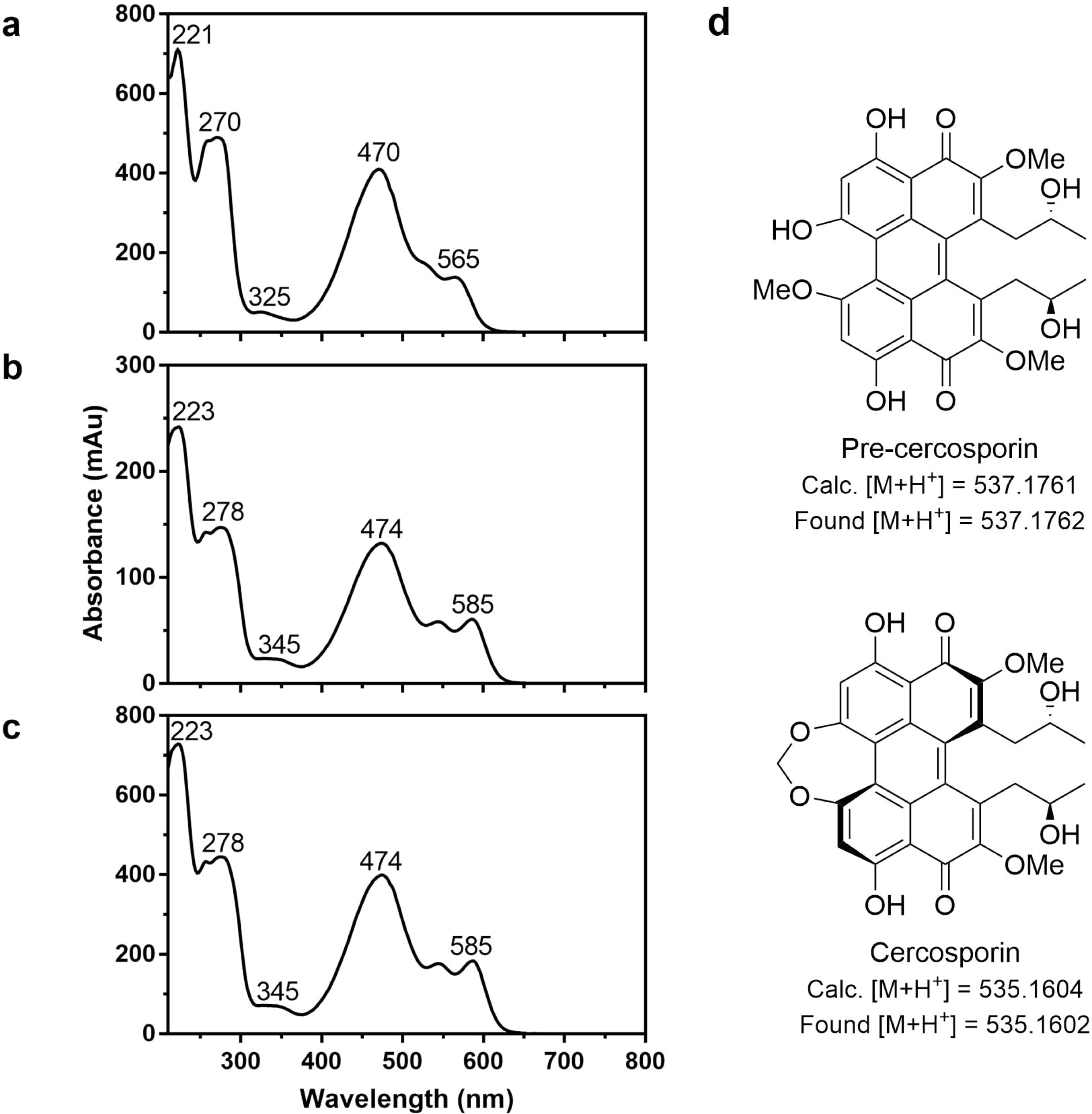
Comparison of UV-Vis spectra of cercosporin and pre-cercosporin. UV-Vis spectra were extracted from 280 nm HPLC chromatograms. Wavelengths of relevant UV maxima are indicated. **a)** 7.25 min. peak (cercosporin) from wild-type *C. beticola*. **b)** 5.36 min. peak (pre-cercosporin) from *C. beticola* ΔCTB9. **c)** UV-Vis spectrum of 5.36 min. peak (pre-cercosporin) from *C. beticola* ΔCTB10. **d)** Proposed structures of cercosporin and pre-cercosporin.

To firmly support the tentative structure of pre-cercosporin, the crude extract of Δ*CTB9* was further purified by reverse-phase HPLC. To obtain sufficient material for ^1^H-NMR analysis, extractions were performed quickly and in the dark to prevent apparent polymerization of pre-cercosporin. The relative instability of pre-cercosporin compared to cercosporin suggests a possible role for the methylenedioxy bridge in overall stability. Immediately evident in the ^1^H-NMR spectrum was the absence of the methylenedioxy singlet at δ5.74 diagnostic of cercosporin, but the appearance of a new methoxyl signal at δ4.28 and a phenol at δ9.25. Consistent with the new asymmetry in pre-cercosporin, two strongly hydrogen-bonded *peri*-hydroxy groups could be seen far downfield at *ca*. 15 ppm and two aryl hydrogens were observed at δ6.92 and δ6.87. That these latter resonances are observed only in pairs, as are the two side chain methyl doublets at *ca*. 0.6 ppm, and the doubling of other signals imply that pre-cercosporin is formed as a single atropisomer having a helical configuration likely identical to that of cercosporin, although it is conceivable CTB9 or CTB10 sets the final stereochemistry.

### Identification of cercosporin from *Co. fioriniae*

Since our phylogenomic analyses suggested that several *Colletotrichum* spp. harbored *CTB* clusters (Figs. 2, 3), we questioned whether any *Colletotrichum* spp. can produce cercosporin. To initially assess this, two *Co. fioriniae* strains (HC89 and HC91) isolated from apple were assayed for cercosporin production using the KOH assay (30). No cercosporin-like pigment was observed in the media under the same conditions that stimulate cercosporin production in *C. beticola*. Since epigenetic modifiers have been used to induce production of SMs in fungal species (31, 32), we questioned whether this strategy could be used to induce cercosporin production in *Co. fioriniae*. Media amended with the histone deacetylase inhibitor trichostatin A (31) induced production of a red cercosporin-like compound into the media. To characterize this red metabolite, mycelia from both *Co. fioriniae* strains were extracted with ethyl acetate. Reverse-phase HPLC analysis as before revealed a peak with a retention time and UV-vis spectrum consistent with cercosporin in both extracts (Fig. 7*A*, *B*). The presence of cercosporin was confirmed by UPLC-ESI-MS (Fig. 7*C*).

**Figure 7.**
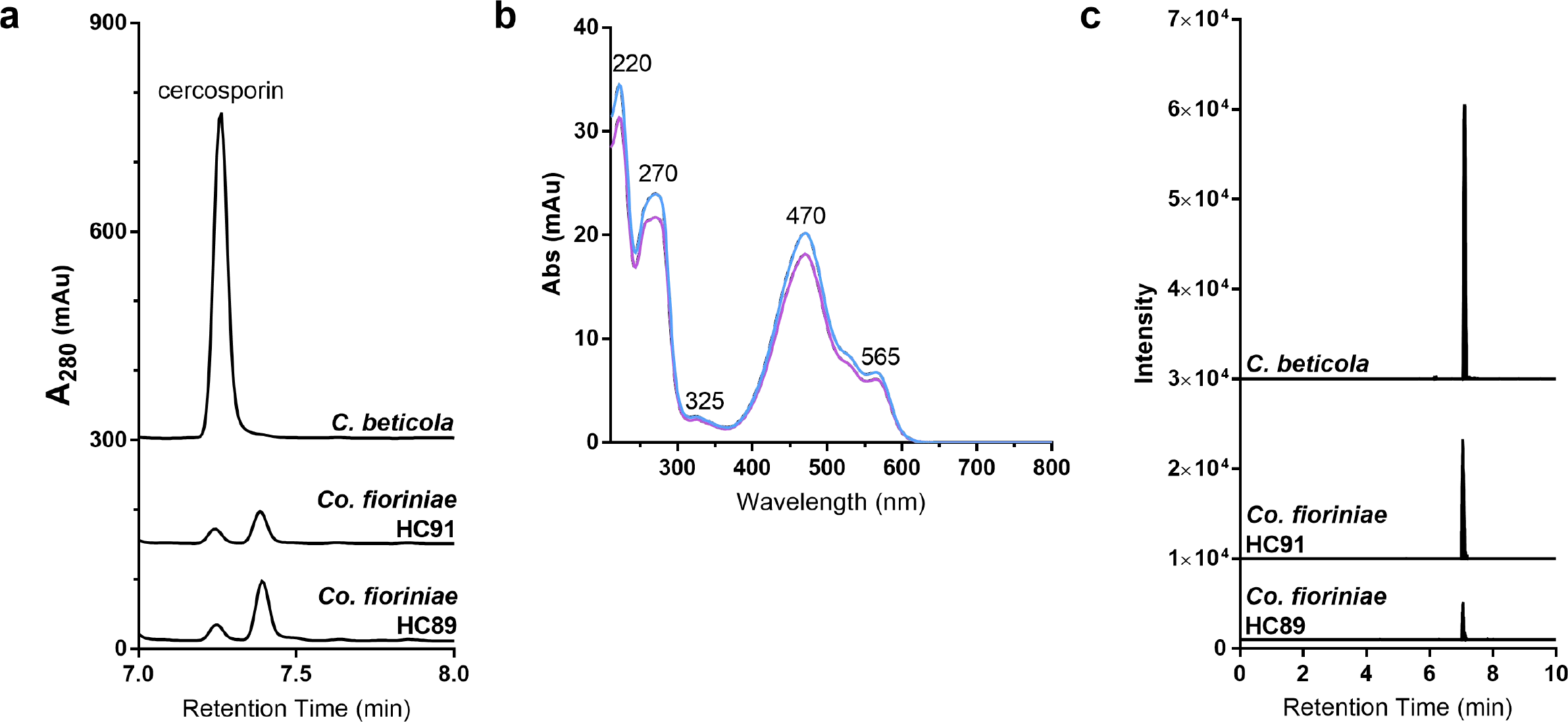
HPLC and UPLC-ESI-MS analysis of *Colletotrichum fioriniae* strains. **a)** HPLC chromatograms at 280 nm of wild-type *C. beticola* and *Co. fioriniae* HC89 and HC91. **b)** UV-Vis spectra of cercosporin (7.25 min. retention time) extracted from *Co. fioriniae* HC89 (blue) and HC91 (purple). Wavelengths of relevant UV maxima are indicated. **c)** Extracted ion chromatograms (m/z = 535.1604) obtained by UPLC ESI-MS, demonstrating cercosporin production in *C. beticola* and *Co. fioriniae* strains HC89 and HC91.

## Discussion

Several hypotheses exist for the maintenance of SM biosynthetic genes as clusters. In one, unlinked SM pathway genes are at a greater risk for dissociation during meiotic recombination (33) or chromosomal rearrangements (34). Additionally, clustering may facilitate strict coordination of gene expression, which may be particularly important during the biosynthesis of SMs that have potentially toxic intermediates to ensure their efficient conversion to final end products (35). Horizontal transfer and maintenance of the ancient *CTB* cluster specifically among plant pathogens suggests that it was critical for disease development in diverse pathosystems, including rice blast caused by *M. oryzae* and various anthracnose diseases caused by *Colletotrichum* spp. on many different crops. The *CTB* clusters in *Co. higginsianum* and *Co. graminicola* were reported as one of the few SM clusters between these species that are microsyntenic (36). Moreover, O’Connell detected specific upregulation of the *CTB* cluster in *Co. higginsianum* during colonization of Arabidopsis (36). Indeed, nine of 14 *Co. higginsianum CTB* genes were among the top 100 most highly expressed genes *in planta*. Recent analysis of natural selection processes in *Co. graminicola* identified orthologs of *CTB* genes *CTB1* and *CFP* as one of ~80 genes undergoing significant positive selection (37), further suggesting a role in pathogenicity. Interestingly, the *CTB* clusters of *Colletotrichum* spp. and *M. oryzae* contain additional genes (two short-chain dehydrogenases, an additional desaturase, a ferric-chelate reductase and an NmrA-like family protein) that have been reported (38) to act as negative transcriptional regulators.

The identification of cercosporin production in two isolates of *Co. fioriniae* has significant implications for the apple packing, storage, and processing industries. Bitter rot, caused by *Colletotrichum* spp., is one of the top pre- and postharvest pathogens of apple (39). This disease is a major problem for the apple industry as it limits fresh fruit in the field and during storage, and has a quiescent stage allowing decay to occur on seemingly high quality apples, only to come out of storage rotten (Jurick II, personal observation). Hence, contamination of processed apple products with cercosporin could be a significant health hazard. For example, other fungal-produced toxins (e.g. patulin, citrinin, penicillic acid) can contaminate processed apple products (40). Patulin, produced by *Penicillium* spp., is the most troubling as it is carcinogenic and consequently the United States and Europe have strict patulin limits in fruit juices and processed pome fruit products (40, 41). Future studies will focus on the role of cercosporin production during the *Colletotrichum*-apple fruit interactions in addition to assaying processed fruit products made from apples with bitter rot symptoms to determine levels of the toxin in fruit. Although only *Co. fioriniae* strains were analyzed for the ability to produce cercosporin, the identification of highly similar *CTB* clusters in other *Colletotrichum* species (Figs. 2, 4) suggest that cercosporin production is wide-spread in this genus.

The microsynteny outside of the established *CTB* cluster prompted us to test whether the flanking genes in *C. beticola* are also required for cercosporin biosynthesis. Notably, we observed that these flanking genes, similar to the established *CTB* genes, were up-regulated under cercospor-ininducing conditions. Furthermore, targeted gene replacement of *CbCTB9, CbCTB10, CbCTB11* and *CbCTB12* completely abolished cercosporin biosynthesis, while replacement of *CbCTB9* and *CbCTB10* also resulted in the accumulation of a new, red metabolite, pre-cercosporin. We thus conclude that the *CTB* cluster is significantly larger than previously described (12).

The isolation and characterization of a new intermediate in the cercosporin biosynthetic pathway, pre-cercosporin, strongly suggests that formation of the unique 7-membered methylenedioxy bridge in the final product is the result of a two-step process requiring three genes. First, one of two precursor aryl methoxyl groups is oxidatively removed (possibly by CTB7, a flavin-dependent oxidoreductase), followed by oxidative ring closure by CTB9, an apparent α-ketoglutarate-dependent dioxygenase, in collaboration with CTB10. The precise role of CTB10, a putative dehydratase, in ring closure is unclear, but it could serve to facilitate closure of the unfavorable 7-membered methylenedioxy ring. In contrast, a single cytochrome P450 is known to convert two aryl *ortho*-methoxyl groups into the relatively more common 5-membered methylenedioxy group in alkaloid biosynthesis (42). A tentative cercosporin biosynthesis scheme was recently proposed (14) without knowledge of the expanded *CTB* cluster. However, in light of the identification of pre-cercosporin and the potential functions of the other newly discovered *CTB* genes, the previously proposed biosynthetic pathway (14) will have to be revised. While these investigations will be reported in due course, we suspect the newly discovered laccase (CTB12) acts early in the pathway to dimerize the product of CTB3 (14) to the first perylenequinone intermediate, which would have precedent in synthetic chemistry (43).

Despite sustained research on cercosporin for several decades, there are significant knowledge gaps in cercosporin biosynthesis. Our data shed new light on cercosporin biology that will have significant impact on cercosporin research specifically and perylenequinone research in general. The finding that at least one species in the important plant pathogenic genus *Colletotrichum* can produce cercosporin has significant implications for disease management. Moreover, since *Co. fioriniae* may secrete cercosporin into apple food products that may be directly consumed by humans, the toxic effects of cercosporin on human health may need to be considered.

## Materials and Methods

For further information, see *SI Appendix* Materials and Methods.

### *Cercospora* and *Colletotrichum* spp. genome sequencing

Genomic DNA of *C. beticola* strain 09-40 was isolated using the CTAB method from mycelia scraped from the surface of V8 juice agar Petri plates (44). Library preparation of three genomic libraries with increasing insert size (500 bp, 5 Kbp and 10 Kbp) and subsequent paired-end (PE) and mate-pair (MP) genome sequencing was performed by BGI Americas Corporation (BGI) using the Illumina platform. For detailed information on *C. beticola* genome assembly, see *SI Appendix* Materials and Methods.

*Cercospora berteroae* strain CBS 538.71 was obtained from Centraal Bureau voor Schimmelcultures (CBS) and cultivated on Petri plates containing potato dextrose agar (PDA; Difco). High-quality DNA was extracted using the CTAB method (45). Library preparation (500 bp) and subsequent paired-end (PE) genome sequencing was performed by BGI via the Illumina platform. A total of 31 million high-quality filtered sequence reads with an average length of 100 bp were generated. A draft genome assembly was constructed using SOAPdenovo (version 2.04), applying default parameters and K-mer length 51.

*Colletotrichum fioriniae* strains HC89 and HC91 were isolated previously from infected apples and cultivated on Petri plates containing potato dextrose agar (PDA; Difco). High-quality DNA was extracted using the CTAB method (45). Library preparation (500 bp) and subsequent paired-end (PE) genome sequencing was performed by BGI via the Illumina platform. Draft genome assemblies were constructed using SOAPdenovo (version 2.04), applying default parameters and K-mer length 51.

### Protein function characterization

For functional characterization of the predicted protein sequences, hardware-accelerated BLASTp on a DeCypher machine (TimeLogic; Carlsbad, USA) was used to identify homologous proteins in the non-redundant (nr) protein database obtained at the NCBI. InterProScan (version 5.44; http://www.ebi.ac.uk/Tools/pfa/iprscan5/) was used to identify conserved protein domains. The results of both analyses were imported into Blast2GO (46) and used to generate single, uniquely functional annotations for each protein as well as a list of all associated gene ontology (GO) terms.

### Secondary metabolite cluster identification, characterization and visualization

Putative SM clusters were identified in the genome sequence of *C. beticola* and that of related fungi using antiSMASH2 (18) (version 2.1.0; https://bitbucket.org/antismash/antismash2/). To generate antiSMASH2-required EMBL formatted genome files, the GFF3 gene features files in combination with the respective genome sequences were converted to the EMBL sequence format using the custom Perl script *GFF3_2_EMBL.pl*. Subsequently, antiSMASH2 was run with default parameters, allowing for the identification of PKS, NRPS SM, Hybrid PKS-NRPS, terpene cyclase (TC), siderophore and lantipeptide SM clusters. SM clusters that showed similarity to a mixture of these clusters or only a minimal set of homologous protein domains were depicted as “other.” In addition, DMAT, for dimethylallyl tryptophan synthase clusters were identified by screening the InterProScan results for Pfam domain PF11991.

### Secondary metabolite phylogenetic analyses

For phylogenetic analyses of the type I polyketide enzymes, we used Mafft (version 7.187), applying global alignment (–globalpair) and a 1000 cycles of iterative refinement (–maxiterate 1000), to align full-length sequences as well as selected domains of all PKS enzymes that were identified by antiSMASH2 in the genome sequences of the six Dothideomycetes: *C. beticola*, *D. septosporum*, *Z. tritici*, *L. maculans*, *P. tritici-repentis* and *P. nodorum*, and one Eurotiomycete: *Aspergillus nidulans*. In addition, previously characterized polyketide synthases (*SI Appendix*, Table S8) were included for reference. Prior to phylogenetic tree reconstruction, the alignments were trimmed with TrimAl (47) (version 1.2). Maximum likelihood phylogenetic trees were determined with RaxML (version 8.1.3), applying rapid bootstrapping (-# 100) and automated protein model selection (-m PROTGAMMAAUTO). Final trees were prepared online using EvolView (48). Species tree topologies were built with Cvtree (49) webserver by uploading the predicted proteomes of 48 published Ascomycete fungi (*SI Appendix*, Table S4).

For phylogenetic tree reconciliation analyses of the protein and species trees, the protein trees were pre-processed with treefixDTL (50) (version 1.1.10) to minimize errors introduced during tree reconstruction. TreefixDTL can correct phylogenetic trees in the presence of horizontal gene transfer. Reconciliation analyses as well as rooting were conducted in NOTUNG (23) according to the instructions (version 2.8 beta).

### Genome-wide gene cluster microsynteny and protein identity analysis

Genome-wide gene-by-gene cluster analyses were performed using the custom Perl script *calClusterSimilarity.pl*, and plotted using ggplot2 in R using *synteny.R*. As input, this pipeline takes the typical output of an orthoMCL analysis, reformatted by *analyseOrthoMCL.pl*. In short, it requires each proteinId to have an associated clusterId. Furthermore, it requires properly formatted GFF3 files for each genome that are used to associate location of protein-coding genes and their flanks. Finally, the number of flanking genes to be used can be chosen freely, but must be set ODD. For the analyses presented in Figure 4, a cluster size of 30 was set. Genome-wide protein-by-protein best-BLAST percent identities were derived from the similarities table prepared during orthoMCL analyses and subsequently plotted in R using *pairwise_pident_boxplots.R*.

### Gene expression analysis

To investigate the expression of cercosporin cluster genes, *C. beticola* was grown in a 250 mL Erlenmeyer flask containing 100 mL potato dextrose broth (PDB; Difco) either in the light or dark, to promote and repress cercosporin production, respectively. Total RNA was isolated using TRIzol (ThermoFisher) following the manufacturer’s instructions followed by an on-column Dnase treatment (Qiagen). Total RNA was used for cDNA synthesis using an oligo-(dT) primer and the SuperScript II reverse transcriptase kit (Invitrogen) following manufacturer’s instructions. The resulting cDNA was used as a template for quantitative polymerase chain reaction (qPCR). Selected genes were queried for expression using the Power SYBR Green PCR Master Mix (Applied Biosytems) using a PTC-2000 thermal cycler (MJ Research) outfitted with a Chromo4 Real-Time PCR Detector (Bio-Rad) and MJ Opticon Monitor analysis software version 3.1 (Bio-Rad). Primers for gene expression analysis are listed in *SI Appendix*, Table S9).

### Transformation and disruption of target genes

Split-marker PCR constructs for targeted gene replacement were prepared as described (44) using genomic DNA of 10-73-4 and 09-40 wild-type *C. beticola* and pDAN as PCR templates. Selected mutants were complemented using pFBT005, which encodes resistance to nourseothricin and allowed us to clone our gene of interest between the ToxA promoter and TrpC terminator using PacI and NotI (Promega) restriction sites. For detailed information, see *SI Appendix* Materials and Methods.

### 4,6,9-trihydroxy-1,12-bis(2-hydroxypropyl)-2,7,11-trimethoxyperylene-3,10-dione (pre-cercosporin)

Data obtained at 5 °C on a Bruker AVANCE spectrometer. ^1^H NMR (400 MHz, CDCl_3_, δ): 15.24 (s, 1H), 14.93 (s, 1H), 9.25 (s, 1H), 6.92 (s, 1H), 6.87 (s, 1H), 4.28 (s, 3H), 4.19, (s, 3H), 4.18 (s, 3H), 3.57 – 3.51 (m, 2H), 3.42 – 3.36 (sym 5-line overlapping signal, 2H), 2.86 – 2.74 (m, 2H), 0.64 (d, *J* = 6.1 Hz, 3H), 0.60 (d, *J* = 6.1 Hz, 3H). UPLC-ESI-HRMS: calculated for C_29_H_29_O_10_ [M+H^+^]: 537.1761, found [M+H^+^]: 537.1762.

### *Colletotrichum* spp. cercosporin assay

To determine whether *Colletotrichum* species were able to produce cercosporin, two monoconidial isolates *C. fioriniae* (HC89 and HC91) were grown on 9 cm Petri plates containing 15 mL of PDA as described above to replicate conditions that were conducive for cercosporin production *in vitro*. Seven day old cultures of each isolate were grown in a temperature controlled incubator at 25 °C with natural light. A pinkish to dark red color was visible in the media for all isolates except HC75 which had a yellow-colored pigment. Using a #2 cork borer, three plugs were removed from each isolate from the edge, middle and center of each colony and placed in small screw cap glass vials. Three plugs were also removed from an uncolonized PDA plate and included as a negative control. Cercosporin (Sigma-Aldrich) was dissolved in acetone to 100 mM and used as a positive control. 5N KOH was added to each vial to cover the surface of the plugs and incubated on a shaking incubator at room temperature for 4 hours. Supernatants were examined for cercosporin spectrophotometrically. To induce cercosporin production, we followed the procedures described by Shwab et al. (31) except 10.0 µM trichostatin A (TSA; Sigma) was used. Cercosporin production by *C. fioriniae* HC89 and HC91 was confirmed by HPLC and UPLC-ESI-MS analysis as described above for pre-cercosporin.

## Acknowledgements

We thank W. Underwood and T. L. Friesen (USDA – ARS) for review of the manuscript, A. G. Newman for helpful discussions and a reference sample of cercosporin and N. Metz (USDA – ARS) for excellent technical assistance. Mention of trade names or commercial products in this publication is solely for the purpose of providing specific information and does not imply recommendation or endorsement by the U.S. Department of Agriculture.

## References

1. Fuckel K (1863) Fungi Rhenani exsiccati, Fasc. I-IV. Hedwigia 2:132–136.

2. Stergiopoulos I, Collemare J, Mehrabi R, De Wit PJGM (2013) Phytotoxic secondary metabolites and peptides produced by plant pathogenic *Dothideomycete* fungi. FEMS Microbiol Rev 37(1):67–93.

3. Goodwin SB, Dunkle LD (2010) Cercosporin production in *Cercospora* and related anamorphs of *Mycosphaerella*. Cercospora leaf spot of sugar beet and related species, eds Lartey RT, Weiland JJ, Panella L, Crous PW, & Windels CE (The American Phytopathological Society), pp 97–108.

4. Daub ME, Ehrenshaft M (2000) The photoactivated *Cercospora* toxin cercosporin: contributions to plant disease and fundamental biology. Annu Rev Phytopathol 38(1):461–490.

5. Daub ME, Hangarter RP (1983) Light-induced production of singlet oxygen and superoxide by the fungal toxin, cercosporin. Plant Physiol 73(3):855–857.

6. Dobrowolski DC, Foote CS (1983) Cercosporin, a singlet oxygen generator. Angew Chem, Int Ed Engl 22(9):720–721.

7. Daub ME (1982) Cercosporin, a photosensitizing toxin from *Cercospora* species. Phytopathol 72(4):370–374.

8. Gunasinghe N, You MP, Cawthray GR, Barbetti MJ (2016) Cercosporin from *Pseudocercosporella capsellae* and its critical role in white leaf spot development. Plant Dis 100(8):1521–1531.

9. Crous PW, et al. (2013) Phylogenetic lineages in *Pseudocercospora*. Stud Mycol 75(1):37–114.

10. Daub ME (1981) Destruction of tobacco cell-membranes by the photosensitizing toxin, cercosporin. Phytopathol 71(8):869–869.

11. Daub ME (1987) Resistance of fungi to the photosensitizing toxin, cercosporin. Phytopathol 77(11):1515–1520.

12. Chen HQ, Lee MH, Daub ME, Chung KR (2007) Molecular analysis of the cercosporin biosynthetic gene cluster in *Cercospora nicotianae*. Mol Microbiol 64(3):755–770.

13. Newman AG, Vagstad AL, Belecki K, Scheerer JR, Townsend CA (2012) Analysis of the cercosporin polyketide synthase CTB1 reveals a new fungal thioesterase function. Chem Commun 48(96):11772–11774.

14. Newman AG, Townsend CA (2016) Molecular characterization of the cercosporin biosynthetic pathway in the fungal plant pathogen *Cercospora nicotianae*. J Am Chem Soc 138(12):4219–4228.

15. Choquer M, Lee MH, Bau HJ, Chung KR (2007) Deletion of a MFS transporter-like gene in *Cercospora nicotianae* reduces cercosporin toxin accumulation and fungal virulence. FEBS Lett 581(3):489–494.

16. Callahan TM, Rose MS, Meade MJ, Ehrenshaft M, Upchurch RG (1999) CFP, the putative cercosporin transporter of *Cercospora kikuchii*, is required for wild type cercosporin production, resistance, and virulence on soybean. Mol Plant-Microbe Interact 12(10):901–910.

17. Chand R, et al. (2015) Draft genome sequence of *Cercospora canescens*: a leaf spot causing pathogen. Curr Sci 109(11):2103–2110.

18. Blin K, et al. (2013) antiSMASH 2.0—a versatile platform for genome mining of secondary metabolite producers. Nucleic Acids Res 41(W1):W204–W212.

19. Collemare J, et al. (2014) Secondary metabolism and biotrophic lifestyle in the tomato pathogen *Cladosporium fulvum*. PLoS ONE 9(1):e85877.

20. Medema MH, Cimermancic P, Sali A, Takano E, Fischbach MA (2014) A systematic computational analysis of biosynthetic gene cluster evolution: lessons for engineering biosynthesis. PLoS Computational Biology 10(12):e1004016.

21. Chooi Y-H, Muria-Gonzalez MJ, Solomon PS (2014) A genome-wide survey of the secondary metabolite biosynthesis genes in the wheat pathogen *Parastagonospora nodorum*. Mycology 5(3):192–206.

22. Koczyk G, Dawidziuk A, Popiel D (2015) The distant siblings — A phylogenomic roadmap illuminates the origins of extant diversity in fungal aromatic polyketide biosynthesis. Genome Biol Evol 7(11):3132–3154.

23. Stolzer M, et al. (2012) Inferring duplications, losses, transfers and incomplete lineage sorting with nonbinary species trees. Bioinformatics 28(18):i409–i415.

24. Tudzynski B, et al. (2003) Characterization of the final two genes of the gibberellin biosynthesis gene cluster of *Gibberella fujikuroi*: *des* and *P450-3* encode GA4 desaturase and the 13-hydroxylase, respectively. J Biol Chem 278(31):28635–28643.

25. Kim J-E, et al. (2005) Putative polyketide synthase and laccase genes for biosynthesis of aurofusarin in *Gibberella zeae*. Appl Environ Microbiol 71(4):1701–1708.

26. Williams JS, Thomas M, Clarke DJ (2005) The gene *stlA* encodes a phenylalanine ammonia-lyase that is involved in the production of a stilbene antibiotic in *Photorhabdus luminescens* TT01. Microbiol 151(8):2543–2550.

27. Choquer M, Lee M-H, Bau H-J, Chung K-R (2007) Deletion of a MFS transporter-like gene in *Cercospora nicotianae* reduces cercosporin toxin accumulation and fungal virulence. FEBS Lett 581(3):489–494.

28. Frandsen RJN, et al. (2011) Two novel classes of enzymes are required for the biosynthesis of aurofusarin in *Fusarium graminearum*. J Biol Chem 286(12):10419–10428.

29. Gao Q, et al. (2011) Genome sequencing and comparative transcriptomics of the model entomopathogenic fungi *Metarhizium anisopliae* and *M. acridum*. PLoS Genet 7(1):e1001264.

30. Choquer M, et al. (2005) The *CTB1* gene encoding a fungal polyketide synthase is required for cercosporin biosynthesis and fungal virulence of *Cercospora nicotianae*. Mol Plant-Microbe Interact 18(5):468–476.

31. Shwab EK, et al. (2007) Histone deacetylase activity regulates chemical diversity in *Aspergillus*. Eukaryot Cell 6(9):1656–1664.

32. Williams RB, Henrikson JC, Hoover AR, Lee AE, Cichewicz RH (2008) Epigenetic remodeling of the fungal secondary metabolome. Org Biomol Chem 6(11):1895–1897.

33. Galazka JM, Freitag M (2014) Variability of chromosome structure in pathogenic fungi - of ‘ends and odds’. Curr Opin Microbiol 20(0):19–26.

34. de Jonge R, et al. (2013) Extensive chromosomal reshuffling drives evolution of virulence in an asexual pathogen. Genome Res 23(8):1271–1282.

35. McGary KL, Slot JC, Rokas A (2013) Physical linkage of metabolic genes in fungi is an adaptation against the accumulation of toxic intermediate compounds. Proc Natl Acad Sci U S A 110(28):11481–11486.

36. O’Connell RJ, et al. (2012) Lifestyle transitions in plant pathogenic *Colletotrichum fungi* deciphered by genome and transcriptome analyses. Nat Genet 44(9):1060–1065.

37. Rech GE, Sanz-Martín JM, Anisimova M, Sukno SA, Thon MR (2014) Natural selection on coding and noncoding DNA sequences is associated with virulence genes in a plant pathogenic fungus. Genome Biol Evol 6(9):2368–2379.

38. Stammers DK, et al. (2001) The structure of the negative transcriptional regulator NmrA reveals a structural superfamily which includes the short-chain dehydrogenase/reductases. The EMBO Journal 20(23):6619–6626.

39. Munir M, Amsden B, Dixon E, Vaillancourt L, Gauthier NAW (2016) Characterization of *Colletotrichum* species causing bitter rot of apple in Kentucky orchards. Plant Dis 100(11):2194–2203.

40. Wright SAI (2015) Patulin in food. Curr Opin Food Sci 5:105–109.

41. Puel O, Galtier P, Oswald I (2010) Biosynthesis and toxicological effects of patulin. Toxins 2(4):613.

42. Díaz Chávez ML, Rolf M, Gesell A, Kutchan TM (2011) Characterization of two methylenedioxy bridge-forming cytochrome P450-dependent enzymes of alkaloid formation in the Mexican prickly poppy *Argemone mexicana*. Arch Biochem Biophys 507(1):186–193.

43. Hauser FM, Sengupta D, Corlett SA (1994) Optically active total synthesis of calphostin D. J Org Chem 59(8):1967–1969.

44. Bolton MD, et al. (2016) RNA-sequencing of *Cercospora beticola* DMI-sensitive and -resistant isolates after treatment with tetraconazole identifies common and contrasting pathway induction. Fungal Genet Biol 92:1–13.

45. Stewart C, Via LE (1993) A rapid CTAB DNA isolation technique useful for RAPD fingerprinting and other PCR applications. BioTechniques 14(5):748–749.

46. Conesa A, Götz S (2008) Blast2GO: a comprehensive suite for functional analysis in plant genomics. Int J Plant Genomics 2008:12.

47. Capella-Gutiérrez S, Silla-Martínez JM, Gabaldón T (2009) trimAl: a tool for automated alignment trimming in large-scale phylogenetic analyses. Bioinformatics 25(15):1972–1973.

48. Zhang H, Gao S, Lercher MJ, Hu S, Chen W-H (2012) EvolView, an online tool for visualizing, annotating and managing phylogenetic trees. Nucleic Acids Res 40(W1):W569–W572.

49. Qi J, Luo H, Hao B (2004) CVTree: a phylogenetic tree reconstruction tool based on whole genomes. Nucleic Acids Res 32(suppl 2):W45–W47.

50. Bansal MS, Wu Y-C, Alm EJ, Kellis M (2014) Improved gene tree error correction in the presence of horizontal gene transfer. Bioinformatics 31:1211–1218.

